# A distinct fingerprint of inflammatory mediators and miRNAs in *Plasmodium vivax* severe thrombocytopenia

**DOI:** 10.1101/2020.08.20.260463

**Authors:** Marina L. S. Santos, Roney S. Coimbra, Tais N. Sousa, Luiz F. F. Guimarães, Matheus S. Gomes, Laurence R. Amaral, Dhelio B. Pereira, Cor J. F. Fontes, Ibrahim Hawwari, Bernardo S. Franklin, Luzia H. Carvalho

## Abstract

**Background:** Severe thrombocytopenia can be a determinant factor in the morbidity of *Plasmodium vivax* (*Pv*), the most widespread human malaria. Although immune mechanisms may drive *Pv*-induced severe thrombocytopenia (PvST), the current data on the cytokine landscape in PvST is scarce, and often conflicting. The analysis of the bidirectional circuit of inflammatory mediators and miRNAs would lead to a better understanding of the mechanisms underlying PvST.

**Methods:** We combined Luminex proteomics, NanoString miRNA quantification, and machine learning, to evaluate an extensive array of plasma mediators in uncomplicated *Pv* patients, whose blood platelet counts varied from reference values to PvST.

**Results:** Unsupervised clustering analysis identified PvST-linked signatures comprised of both inflammatory (CXCL10, CCL4, and IL-18) and regulatory (IL-10, IL-1Ra, HGF) mediators. As part of PvST signatures, IL-6 and IL-8 were critical to discriminate *Pv* subgroups, while CCL2 and IFN-γ from healthy controls. Supervised machine learning spotlighted IL-10 in *Pv*-mediated thrombocytopenia, and provided evidence for a potential signaling route involving IL-8 and HGF. Finally, we identified a set of miRNAs capable of modulating these signaling pathways.

**Conclusions:** The results place IL-10 and IL-8/HGF in the center of PvST and propose investigating these signaling pathways across the spectrum of malaria infections.

## Introduction

*Plasmodium vivax* is the most widespread of the human malaria parasites, placing 3.3 billion people at risk worldwide [1]. More recently, *P. vivax* (*Pv*) burden has been aggravated by growing evidence of its presence across all regions of Africa [2]. Challenges to the control and elimination of *Pv* include its ability to relapse, its remarkable transmission efficiency, and low-density blood-stage infections, often undetected by routine surveillance [3].

It is currently a consensus that the virulence of *Pv* has been underestimated [4–6], particularly in the presence of co-morbidities [7]. While there are critical gaps in the current knowledge of *Pv* pathophysiology, it is well-established that vivax malaria is associated with a systemic inflammatory response [8], perhaps more intense than its counterpart *P. falciparum* [7, 9], which is more commonly associated with severe malaria. Findings suggest that tissue accumulation of *Pv* may occur, with the hidden biomass greatest in severe disease and capable of mediating systemic inflammatory response [10][11].

*Plasmodium vivax*-induced severe thrombocytopenia (PvST), characterized by blood platelet counts below 50,000 per mm^3^, is a common clinical complication in *Pv* malaria [12–14]. The mechanisms leading to PvST are unclear but may be related to platelet activation, consumption and/or phagocytosis [15] [16] [17].

Growing evidence strengthen an essential role of platelets as mediators of inflammation through their capacity to secrete numerous proteins upon activation or via their interaction with the endothelium, or with leukocytes [18]. For example, we have recently demonstrated that platelets enhance the inflammasome activity of innate immune cells and amplify IL-1-driven inflammation [19]. Therefore, it is possible to speculate that platelets may be critical players in *Pv*-mediated systemic inflammatory response.

The role of platelets in malaria is complex and multifaceted [20]. While platelets can kill circulating parasites of all major human *Plasmodium* species through the release of platelet factor 4 (PF4 or CXCL4) [21], most studies indicate a predominantly deleterious role [22], which may involve the von Willebrand factor [23], the coagulation cascade [24], and the protein C pathway [25]. Additionally, studies involving controlled human *Pv* malaria infection (CHMI) suggested a link between platelets and endothelial activation, an essential pathogenic process in severe malaria [26].

Given the multifactorial mechanisms through which platelets can impact *Pv* malaria, we investigated here crucial factors and pathways that could underlie a fingerprint of PvST. For that, we measured the plasma concentrations of cytokines, chemokines, and growth factors in a cohort of *P. vivax* patients with varying degrees of thrombocytopenia. To gain additional insights into possibly perturbed regulatory pathways in PvST, we included a group of microRNAs (miRNAs), a class of small non-coding RNAs that regulate gene expression and seem to be critical to regulate platelet function [27]. By combining these highly sensitive methods with machine learning algorithms, we provide here essential insights into the interplay between inflammatory mediators, miRNAs, and *P. vivax*-induced severe thrombocytopenia.

## Patients and Methods

### Study participants and sample collection

Individuals who sought care at Brazilian malaria reference healthcare facilities in endemic areas of the Amazon Region and presented *Pv*-positive thick blood smear were invited to participate in the study. Exclusion criteria consisted of: (i) refusal to provide written informed consent; (ii) age below 17 years; (iii) self-reported pregnancy; (iv) mixed malaria infections (PCR-based assays); and (v) any other traceable co-morbidities. Upon enrolment, we used a standardized questionnaire to record demographical, epidemiological, clinical and hematological data. Seventy-seven symptomatic uncomplicated *Pv* patients, with a median age of 39 years and a proportion male: female of 4.5: 1, were enrolled in the study (Supplementary Table 1). The interquartile range of parasitaemia was 1,900 to 7,380 parasites per mm^3^, with anaemia present in 25 (32%) of *Pv* patients. Fifty-four (70%) patients present thrombocytopenia (i.e., platelets below150,000/mm^3^), with 9 (12%) of them classified as severe thrombocytopenia (platelets below 50,000/mm^3^), with no evidence of bleeding. Peripheral blood sample (5 mL in EDTA) was collected for each individual; plasma samples were immediately obtained after blood sampling (1,500 × g for 15 min at room temperature) and stored at −80°C until use. Additionally, plasma samples from nine age-matched non-infected healthy adult from the same endemic areas were collected as described above.

The methodological aspects of this study were approved by the Ethical Committee of Research on Human Beings from the René Rachou Institute – Fiocruz Minas (protocols # 05/2008 and # 80235017.4.0000.5091), according to the Brazilian National Council of Health. The study participants were informed about the aims and procedures and agreed with voluntary participation through written informed consent.

### Multiplex determination of inflammatory mediators

The plasma concentrations of 45 cytokines/chemokines/growth factors were measured with the ProcartaPlex^®^ 45-Plex Human array (eBioscience, USA) using Luminex^®^ xMAP technology (MAGPIX, Thermo Fisher Scientific, USA), as recommended.

### Plasma RNA extraction and confirmation by real time PCR

Circulating and exosomal RNA were purified from 400 μL of plasma using Plasma/Serum Circulating and Exosomal RNA Purification kit (Norgen Biotek Corp., Canada), according to manufacturer instructions. In order to control extraction efficiency, a panel contained 5 spiked in probes (ath-miR-159a, cel-miR-248, cel-miR-254, osa-miR-414, osa-miR-442; IDT Technologies, USA) was added after second lysis buffer incubation. After purification, RNA portion was concentrated using RNA Clean-Up and Concentration kit (Norgen Biotek Corp., Canada), following manufacturer instructions. As a control of the RNA extraction, samples were amplified for the U6 (RNU6‑ 1) snRNA by using TaqMan^®^ MicroRNA Reverse Transcription kit, followed by qPCR amplification using QuantStudio 6 Flex (Applied Biosystems, USA), according to fabricant’s protocol. The U6 snRNA was successfully detected (data not shown).

### Analysis of microRNA (miRNA) expression

Detection of the miRNA was performed using NanoString nCounter technology (USA), according to manufacturer’s protocol. The Human v3 miRNA CodeSet allowed multiplex assessment of 800 miRNAs by specific molecular barcodes. Raw data was analysed with NanoString nSolver 4.0, using default quality control standards. For normalization, spike in oligos (cel-miR-248 and cel-miR-254) were used. Following manufacturer’s instructions, only targets that presented average raw count above 100 were considered for further differential expression analysis in Partek Genomics Suite 7.0. Principal component analysis plot was also generated with the same software with batch effects removed. Targets presenting fold change equal or above 1.25 (p<0.05) were considered differentially expressed.

## Data analysis

Differences in medians were tested using Mann-Whitney test or Kruskal-Wallis test, as appropriated. Correlation between variables was assessed by using either Pearson’s or Spearman’s (rank) correlation (GraphPad Prism 7, San Diego, CA, USA) at significance level of 5%. The ensemble of inflammatory mediators differentially expressed in the study population was identified with the algorithm Comparative Marker Selection (fold change ≥ 1.5, Bonferroni p ≤ 0.05) followed by hierarchical clustering (Spearman’s correlation, average linkage) using the GenePattern platform (Broad Institute, MIT, USA). The independent association between inflammatory mediator and the number of platelets was evaluated by adjusting a negative binomial (NB) regression model with stepwise backward deletion. NB regression analysis was performed using the statistical package Stata version 12 (Stata Corp., Texas, USA). Covariates were selected for inclusion in the regression models if they were associated with the outcome at the 15% level of significance according to the exploratory unadjusted analysis. Associations with p values < 0.05 were considered significant. Decision trees were used to select the minimal set of phenotypic features that efficiently segregated groups. The J48 method, present in Weka software (Waikato Environment for Knowledge Analysis, version 3.6.11, University of Waikato, New Zealand), was used for decision tree construction. The Leave-one-out cross-validation (LOOCV) was calculated to estimate the accuracy of the generated model. Functional enrichment analysis of the miRNAs targets with experimental evidences was performed with Ingenuity Pathway Analysis (IPA) (Qiagen) with default parameters.

## Results

### A cytokine/chemokine signature for P. vivax-induced severe thrombocytopenia

To acquire a reliable assessment of the landscape of circulating cytokines/chemokines in P. vivax malaria, we used high throughput Luminex cytokine arrays to profile the concentrations of 45 human proteins (cytokines, chemokines, and growth factors) in the plasma of P. vivax patients (Supplementary Table 1). As baseline comparison, we included plasma from nine age-matched healthy volunteers from the same localities. We found significantly altered plasma concentrations of 15 proteins in *Pv* patients compared to healthy donors (Supplementary Figure 1A). Notably, several proteins in plasma from *Pv* patients correlated with blood platelet counts (Supplementary Figure 1B, and Figure 1). Except for IL-1β and bNGF, the correlation for all other proteins was negative (Supplementary Figure 1), suggesting an involvement of these proteins in the pathogenesis of *P. vivax* associated with severe thrombocytopenia (PvST).

**Figure 1.**
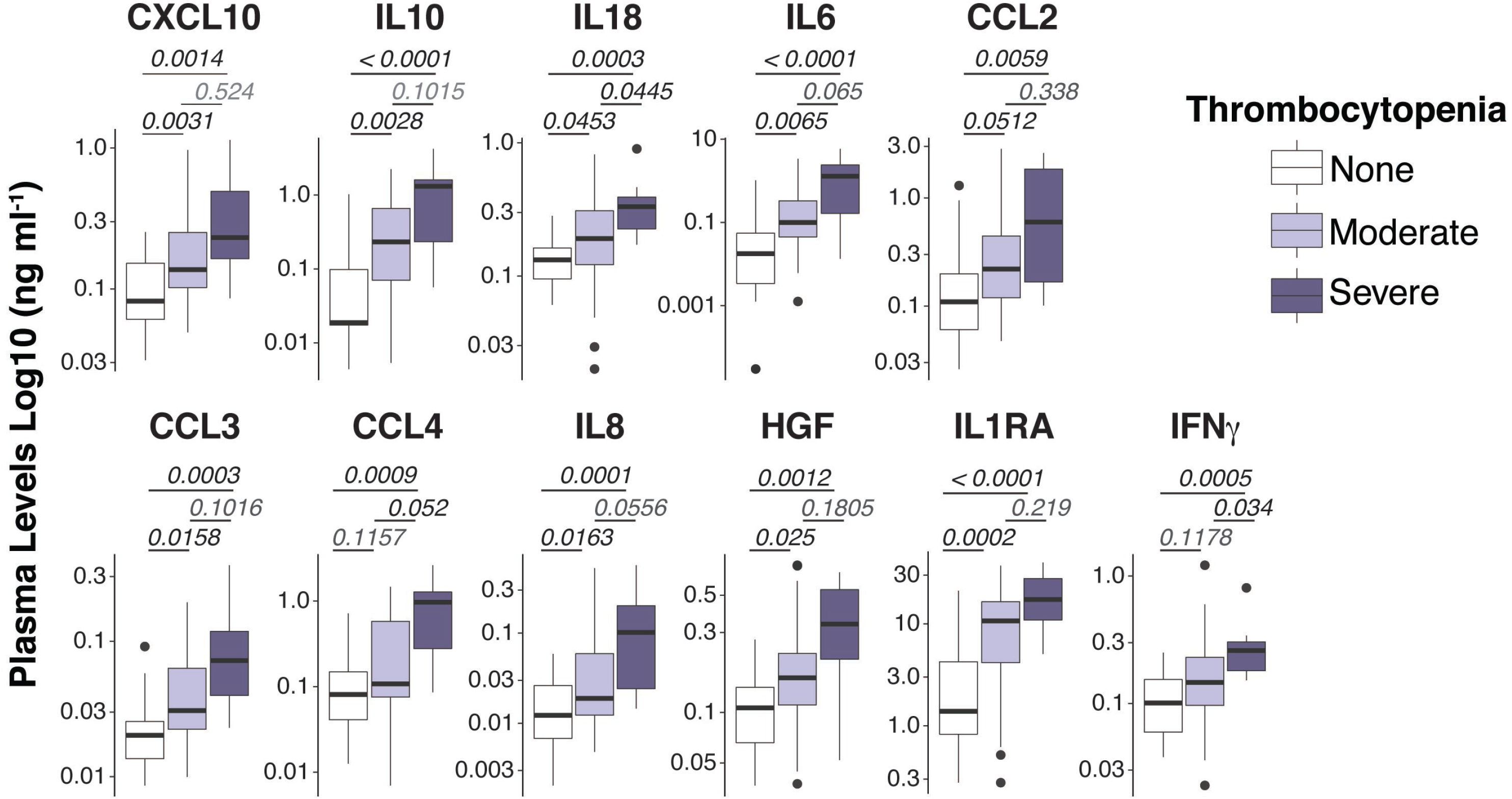
Spectrum of plasma mediators in *P. vivax* infection. Luminex Cytokine plex of cytokines, chemokines and growth factors measured in the plasma of *P. vivax* patients. Patients were stratified according to their blood platelet counts as severe (< 50,000 mm^3^) or moderate (50,000 – 150,000 mm^3^) thrombocytopenia, and non-thrombopenic (≥ 150,000 mm^3^). For each boxplot, median and interquartile ranges were represented as transversal lines in the center and lower/upper bounds of the box, respectively, with whiskers the minimum and maximum values. P values for Kruskal-Wallis multiple comparisons are shown.

To gain additional insights into the relationship between circulating inflammatory mediators and PvST, we applied hierarchical clustering, an unsupervised machine learning algorithm, to investigate patterns of plasma proteins that could reliably report PvST. From all the cytokines that showed correlations with platelet counts (Figure 1 and Supplementary Figure 1B), this analysis highlighted eight proteins (CXCL10, CCL2, CCL4, IL-10, IL-1Ra, IFN-γ, IL-18, and HGF) as part of a specific PvST signature, as compared to unexposed healthy controls (Figure 2A). When we stratified *Pv* patients into subgroups with severe thrombocytopenia or non-thrombocytopenia, this analysis additionally identified IL-8 and IL-6, but not IFN-γ and CCL2 among the eight proteins found in the former comparison (Figure 2B). Using a multivariate regression analysis, we excluded the effect of possible confounding factors, such as *Pv* parasitic density, hemoglobin concentrations and WBC counts. Notably, the following mediators linked to PvST remained independently associated with platelet counts in *Pv* patients: IL-8 (β= −0.0072; p=0.006), IL−10 (β= −0.0009; p=0.005), HGF (β= −0.0026; p<0.001), and CCL2 (β= −0.0005; p= 0.006).

**Figure 2.**
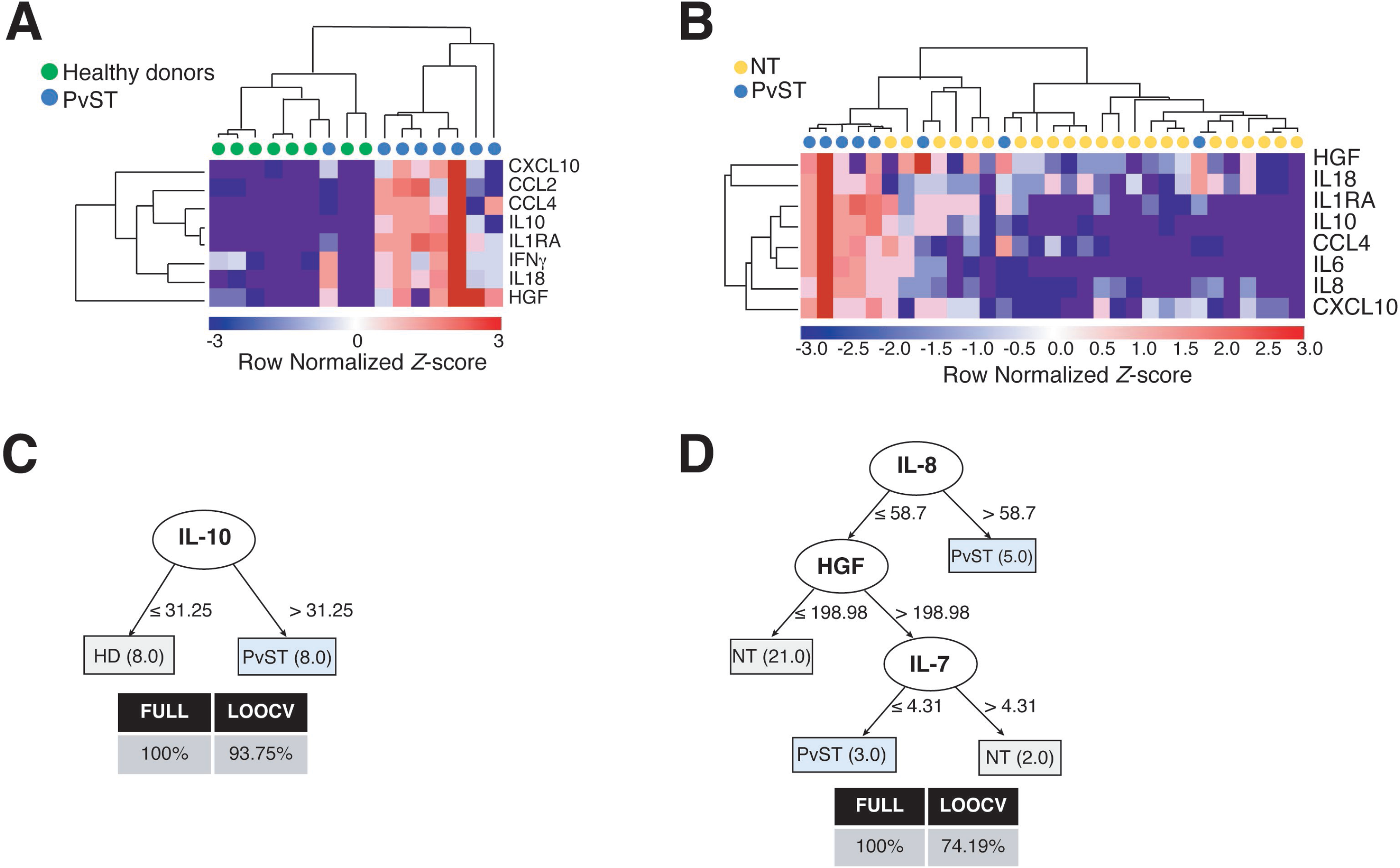
Identifying a cytokine/chemokine signature for *P. vivax* patients with severe thrombocytopenia (PvST). *A-B*, Agglomerative hierarchical clustering showed key cytokines that could distinguish PvST (blue circles) from (*A*) healthy donors (green circles), or (*B*) non-thrombopenic patients (NT, yellow circles). Data were represented as heatmap of clustered proteins (rows) and individual plasma samples (column), with minimum and maximum normalized levels showed in blue and red scales, respectively. Hierarchical clustering was performed based on Spearman’s correlation coefficient, using the average linkage method (GenePattern, Broad Institute). *C-D,* Best-fit decision trees generated with the J48 algorithm (Weka software) identified IL-10 (*C*) and IL-8/HGF/IL7 (*D*) as minimum mediations that efficiently segregated PvST from healthy controls (HD) and from non-trombocytopenic patients (NT), respectively. Weighted of the attribute (plasma levels) were placed in the root of the tree according to the cytokine/chemokine value (pg/mL) that best divided groups. The total of classified registers (correct and incorrect) for each class are given in parentheses for each terminal node with the Full training (FULL) and Leave-one-out cross-validation (LOOCV) accuracies. If incorrectly classified registers exist, they will appear after slash "/".

Next, we employed supervised machine learning to the complete dataset to generate decision trees that could highlight key markers that predict severe thrombocytopenia from possible noise of other mediators. From all inflammatory mediators analyzed, the J48 algorithm generated best-fit decision trees that highlighted (i) IL-10 to discriminate healthy controls from PvST (Figure 2C) and (ii) IL-8, HGF, and IL-7 to discriminate PvST from non-thrombocytopenic patients (Figure 2D). Notably, inclusion of additional biomarkers did not result in an appreciable increase in classification accuracy. Altogether, these findings reveal a panel composed of IL-8/HGF/IL-7 and IL-10 in the center of PvST.

### A signature of miRNAs for *P. vivax*-induced severe thrombocytopenia

MicroRNAs (miRNAs) are well-known regulators of cytokine expression. To gain an additional layer on the mechanisms driving PvST, we used a highly sensitive molecular barcode NanoString approach to examine the miRNA profile of a representative subset of the studied population. The samples were comprised of 26 age-matched *P. vivax* patients, selected based on their blood platelet counts (as none, moderate, or severe thrombocytopenia), as well as healthy individuals, as baseline controls. Malaria patients and healthy individuals clustered into different expression ellipses in the principal component analysis (Figure 3A), which explained 74% of the variation in the dataset. While we detected 17 out of 800 miRNAs in *P. vivax* plasma, only six of them differed between subgroups of *P.vivax* patients or healthy controls (Figure 3B). Within the differentially expressed miRNA between *P. vivax* patients and controls we found that the miRNA pair hsa-miR-4454/hsa-miR-7975 was upregulated (1.957 fold, p = 0.0434) while three other miRNAs were downregulated (< 1.5-fold) in *P. vivax* patients (Figure 3B).

**Figure 3.**
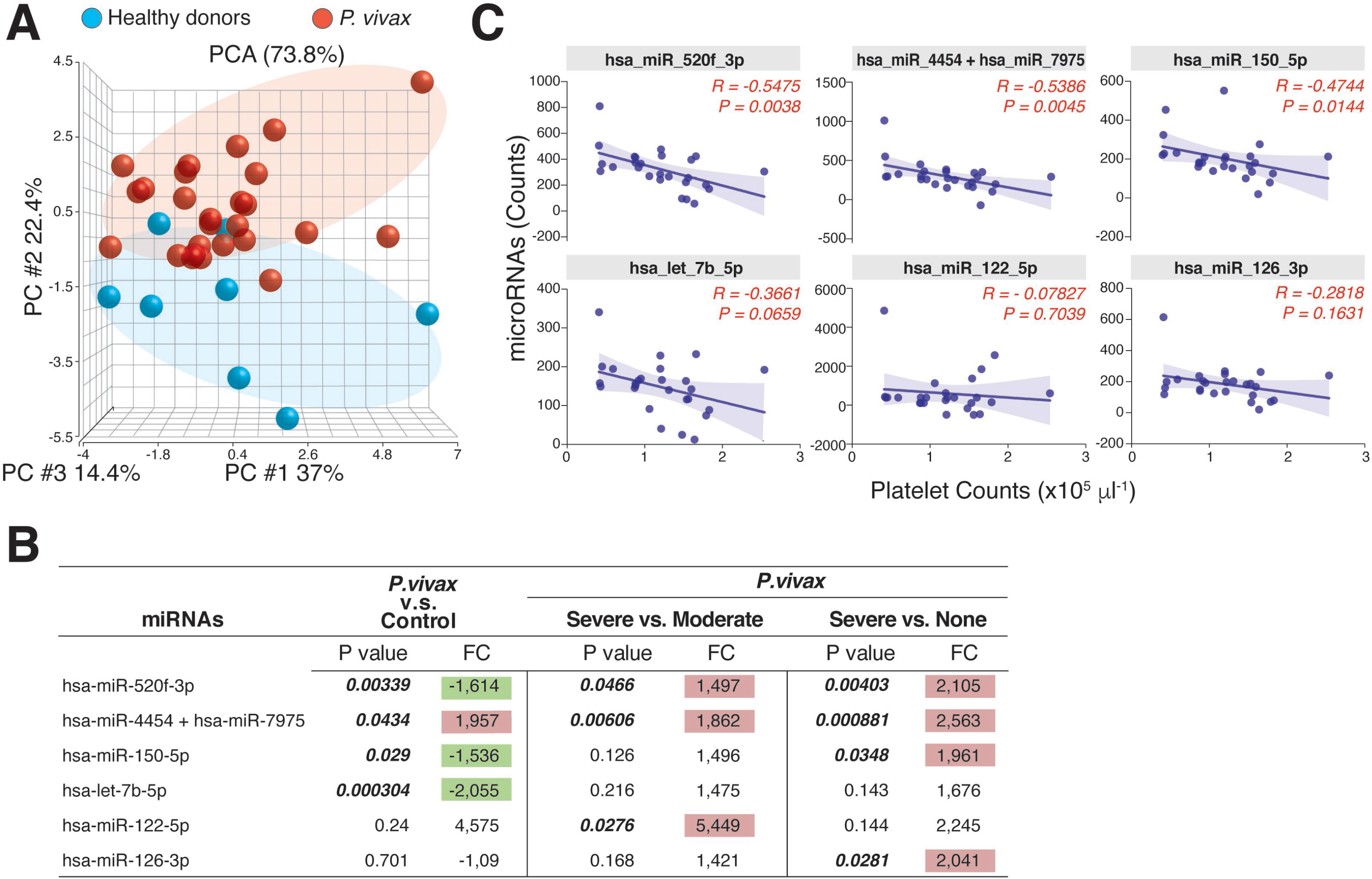
The profile of circulating miRNAs in *P. vivax* patients. *A*, Principal component analysis of the distribution of miRNAs detected in the plasma of a subgroup of P. vivax patients (red, n = 26) and healthy donors (blue, n = 8); *B,* Six miRNAs were differentially expressed in groups of *P. vivax* patients and healthy subjects, with positive values of fold change (FC) highlighted in red and negative values in green, considering p<0.05. *P. vivax* patients were stratified according to their blood platelet counts as: severe (< 50,000 mm^3^) or moderate (50,000 to 150,000 mm^3^) thrombocytopenia, and non-thrombocytopenic patients (≥ 150,000 mm^3^); *C,* Correlations between plasma miRNA levels and blood platelet counts in *P. vivax* patients from *A.* Correlation coefficients and P. values are shown.

The pair hsa-miR-4454/hsa-miR-7975 was additionally significantly increased in patients with severe thrombocytopenia compared to other infections (none, or moderate thrombocytopenia) (Figure 3B). The miRNA hsa-miR-122-5p showed the highest expression levels in PvST patients compared to patients with moderate thrombocytopenia, but this difference was not significant when compared to non-thrombogenic patients. Together, these findings highlight these miRNAs as associated with PvST. Supporting these findings, a correlation between the miRNA levels detected by NanoString and the blood platelet counts in these 26 patients confirmed that platelet counts are negatively associated with plasma levels of miRNAs (Figure 3C). Importantly, plasma miRNA levels did not show significant correlations with other blood parameters (data not shown).

In parallel, the expression of all but one miRNA linked to PvST increased alongside the plasmatic concentration of cytokines, chemokines and growth factor (Supplementary Figure 2), which were highlighted as part of a signature for PvST (Figure 2). Lastly, data analysis of miRNA expression confirmed that the set of miRNAs identified here are involved in key canonical pathways related to the signature of PvST, including the signaling of IL-6, IL-7, IL-8/CXCL8, interferon and IL-10 (Supplementary Table 2). Additionally, signaling pathways related to several stages of the inflammatory process and immune response were also identified.

Finally, the profile of the 17 miRNAs that were detected in plasma samples of the studied subjects generated accurate decision trees based on a single miRNA for severe thrombocytopenia versus healthy controls (has-miR-4454/has-miR-7975) (Figure 4A), or non-thrombocytopenic *P. vivax* infections (has-miR-150-5p) (Figure 4B). Together, our findings provide an intricate relationship between key cytokines/chemokines with their regulatory miRNAs in *P. vivax* malaria that may be useful to define PvST.

**Figure 4.**
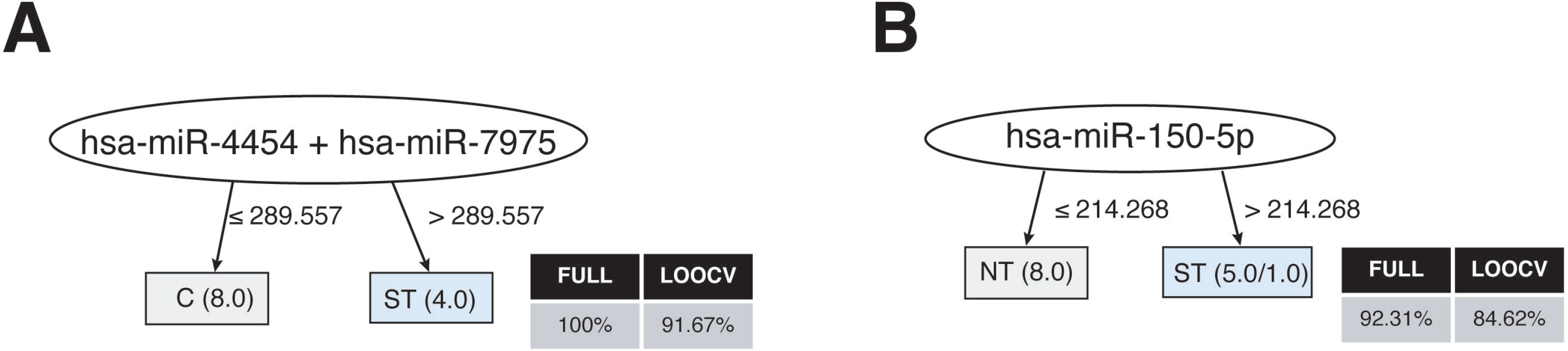
Decision trees for the miRNAs linked to P. vivax severe trombocytopenia. Accurate decision trees generated with the J48 algorithm (Weka software) identified has-miR-4454/has-miR-7975 (*A*) and has-miR-150-5p (*D*) as minimum miRNA that efficiently segregated PvST (ST) from health controls (HC) and from non-trombocytopenic patients (NT), respectively. Weighted of the attribute (miRNA) were placed in the roof of the tree according to the miRNA value (counts) that best divided groups. The total of classified registers (correct and incorrect) for each class are given in parentheses for each terminal node with the Full training (FULL) and Leave-one-out cross-validation (LOOCV) accuracies. If incorrectly classified registers exist, they will appear after slash "/"

## Discussion

Scarce, fragmented, and occasionally conflicting data are available on the cytokine/chemokine network in PvST [14, 16, 28, 29]. Here, we hypothesize that the combined analysis of circulating proteins and miRNAs could provide a more realistic portrait of PvST. We, therefore, examined an extensive array of soluble plasma factors in a cohort of *Pv* patients, whose most prominent hematological alteration was blood platelet counts, which varied from normal ranges (> 150 x10^3^/mm^3^) to severe thrombocytopenia (< 50×10^3^/mm^3^). As expected, no bleeding was reported in our patients, suggesting that megakaryocytes were able to release mega platelets in the circulation to compensate for the low absolute number of platelets in the periphery [15]. In accordance, we found that the mean platelet volume (MPV), a measurement of the platelet size that increases according to platelet destruction, correlated negatively with the platelet counts (r = −0.5426, p = 0.0002).

For the detection of soluble plasma mediators, we used a Luminex-based technology that is highly reproducible compared to conventional cytokine bead arrays [30]. The hierarchical clustering of samples allowed us to define two panels comprised of inflammatory and regulatory mediators that discriminated PvST from health controls or non-thrombocytopenic patients (Supplementary Table 3). Although the same regulatory mediators (IL-10/IL-1Ra/HGF) were included in both panels, decision trees confirmed the levels of IL-10 as critical to classify PvST as compared with controls. Previous studies supported a similar association between IL-10 and decreased blood platelet counts [14, 16, 28, 29].

High levels of IL-10 are commonly observed in *Pv* infections and are primarily associated with the immune system’s effort to counteract excessive inflammation [31]. This oversimplified association of IL-10 with less severe vivax disease has been challenged by studies that reported a lack of correlation between regulatory cytokines and milder symptoms. On the contrary, high levels of IL-10 are linked to intense paroxysms [32], increased disease-severity [33], parasite-related inflammation [10], and the occurrence of recrudescence of blood-stage infections [34]. Although previous studies highlight that IL-10 has complex and not well-characterized functions in *P. vivax* pathogenesis [10], our findings strengthen this cytokine’s critical contribution in *P. vivax*-induced thrombocytopenia.

In addition to IL-10, our analysis identified IL-1Ra as associated with *Pv*-severe thrombocytopenia. IL-1Ra is a naturally occurring member of the IL-1 family that binds to IL-1 receptors and antagonizes IL-1α/IL-1β [35]. Notably, IL-10 is a potent inducer of IL-1Ra, which may represent a mechanism whereby IL-10 exerts its anti-inflammatory effects [36]. Interestingly, the decision tree further indicated HGF as strongly associated with PvST, particularly to differentiate subgroups of *P. vivax* patients. HGF exerts potent anti-inflammatory effects, which seem to involve a signaling cascade leading to increased expression of IL-1Ra [37]. In agreement with the upregulation of IL-1Ra in thrombocytopenic patients, we recently demonstrated that low platelet counts in *Pv* malaria are associated with a progressive decrease in plasma concentrations of IL-1β [19]. While IL-1Ra in *Pv* infections has been little explored, elevated IL-1Ra levels have been associated with increased disease severity in *P. falciparum* malaria-infected children [38]. These findings suggest that an excessive anti-inflammatory response may dampen the necessary inflammatory response able to control the infection [39].

Combined with the abovementioned regulatory mediators, the panels capable of differentiating PvST from other subgroups included well-known inflammatory mediators (Supplementary Table 3). A range of different cell types produces these cytokines/chemokines, which account for the cascade of events that lead to leukocytes recruitment, trafficking, and amplification of inflammation and *Pv* pathogenesis [8, 10, 40]. Interestingly, the up-regulation of IL-6 and IL-8 was critical to discriminate subgroups of vivax patients, but not from healthy subjects. IL-6 has an established role in *Pv* infection, particularly as a marker of systemic inflammation leading to organ dysfunction and disease severity [10]. The same authors demonstrated associations between a decreased activity of plasma ADAMTS13 (a von Willebrand factor cleaving protease), lower platelet counts, and increased concentrations of IL-6, a well-known specific inhibitor of ADAMTS13.

An unexpected finding was the potential involvement of IL-8 in PvST, further validated by the decision trees. IL-8, whose expression is induced by Toll-like receptor (TLR) and IL-1R-stimulated-NF-κB signaling, is a key mediator of neutrophil recruitment [41]. Though IL-8 has been largely underestimated in *P. vivax* infection, a previous study noticed impaired chemotaxis of neutrophils towards an IL-8 gradient, suggesting a possible mechanism for secondary bacterial infection during *Pv* malaria [42]. On the other hand, earlier studies on *P. falciparum* have reported elevated IL-8 levels in patients suffering from severe disease [43, 44]. Although IL-8-mediated thrombocytopenia has not been investigated in malaria, a significant body of evidence suggests its involvement in platelet production, destruction, and/or activation [45–48].

While the mechanism of IL-8 induced PvST is not known, our study identified a potential route involving HGF. Notably, HGF binds to cMet (a receptor tyrosine kinase) to regulate IL-8 expression [49]. HGF itself has been reported to affect the proliferation and differentiation of hemopoietic stem and progenitor cells [50]. Intriguingly, a variant beta-chain of HGF forms a molecular complex with IL-7, and this naturally occurring hybrid cytokine IL-7/HGFβ exerts a potent influence on primitive hematopoietic cells [51]. In the current study, the plasma levels of circulating IL-7 (a growth factor for lymphocytes) were close to the detection limits of the assay (average 4 pg/mL), which precluded definitive conclusions about its involvement in PvST; even though the decision tree algorithm identified IL-7 as useful to classify a small part of *Pv* patients. Despite that, reduced peripheral levels of IL-7 have been associated with inefficient erythropoietic responses in *P. falciparum*-induced severe anemia [52]. Unfortunately, no data related to blood platelet counts were available in the abovementioned pediatric study. In our study, anemia was not a confounding factor as (i) multivariate regression analysis confirmed that hemoglobin levels were not a confounding factor for the association between plasma concentrations of IL-8 and HGF and peripheral platelet counts (ii) the majority of enrolled patients does not present anemia. Collectively, our results suggest that a mechanism involving the upregulation of IL-8 and HGF is involved in PvST. . The potential contribution of the downregulation of IL-7 should be further confirmed.

Considering that cytokines/chemokines are among the most relevant proteins whose expression is regulated by miRNAs [53], we further identify key regulatory miRNAs that could be involved with the inflammatory profile of PvST. Remarkably, a set of six miRNAs were differentiated expressed between PvST patients and other subgroups (non-thrombocytopenic or healthy controls), with the expression levels of all but one miRNA increasing with the circulating levels of strategic mediators such as IL-6, IL-8, IL-10, and HGF.

It is noteworthy that platelet-derived microparticles are transport vehicles for large numbers of miRNAs [54]. Among the miRNAs linked to PvST, we detected platelet-related miRNAs (e.g., miR-126-3p and miR-150-5p), which are known to mediate platelet function and reactivity [54, 55]. This is a relevant observation as we have previously demonstrated that platelets are major sources of circulation microvesicles in *Pv* malaria [56] and that thrombocytopenia strongly correlates with levels of circulating nucleic acids [57]. Lastly, the set of miRNAs identified here were involved in key canonical pathways related to the signature of PvST, including the signaling of IL-6, interferon, IL-8 and IL-10. Interestingly, the same set of miRNA identified the signaling of IL-17A (IL-17, a key component of innate and adaptive immunity) as associated with PvST. Notwithstanding, the plasma levels of IL-17A in our *Pv* patients did not correlate with platelet counts. Perhaps IL-17A plays a more complex role in the cascade of events that lead to PvST. IL-17A signaling mediates the production of chemokines/cytokines such as IL-8, CCL2 and IL-6 [58], which were associated with PvST in our analysis. Altogether, these findings are relevant, as the miRNA profile in *P. vivax* remains poorly explored.

This study has limitations that should be considered when interpreting the results. First, our study included a relatively small number of participants, which may have underpowered some of the statistical analyses. Second, the multivariate model used to control confounding variables was restricted to a small number of participants, which may have excluded cytokines that could be independently associated with platelets. Despite this limitation, key mediators such as IL-10, IL-8, and HGF were clearly identified as independently associated with platelet counts. Third, one single time-point sampling is unlikely to provide insights into the sequence of events from different levels of thrombocytopenia to the progression of clinical disease. Notwithstanding, the direction of the associations with severe thrombocytopenia revealed structural patterns in such data sets that allowed us to identify novel candidate cytokines/chemokine patterns that may be critical for PvST. Notably, our analysis highlighted key core markers such as IL-8 and HGF as potential, and unexplored, signaling routes involving platelets and *Pv* malaria infection. Finally, we provide evidence for an unprecedented role of the HGF signaling in a regulatory pathway involving IL-10/ IL-1Ra. Coherently, this study identified a set of miRNAs capable of modulating these vital chemokine/cytokine pathways, which should be investigated in the context of thrombocytopenia across the spectrum of malaria infections.

## Acknowledgments

We thank malaria patients and healthy controls for participation in the study; the local malaria control team in Porto Velho (RO) and Cuiabá (MT) for their logistic support; Lucas Secchim Ribeiro for fruitful discussions and Cornelia Rohland for sample shipment; the Programme for Technological Development in Tools for Health-PDTIS-FIOCRUZ for use of the Real-Time PCR (RPT09D) facilities; the Coordination for the Improvement of Higher Education Personnel (CAPES) and the Program for Institutional Internationalization of CAPES-PrInt / FIOCRUZ is also acknowledged.

## Notes

## Authorship

Conception and study design: LHC and BSF. Fieldwork and recruitment of participants: MLSS, CJFF, DBP. Performance of experiments: MLSS and IH. Data analysis: MLSS, IH, TNS, CJFF, LHC, BSF. Bioinformatics analysis: RSC, MSG, LRA. The drafting of the manuscript: LHC and BSF. All authors critically revised the manuscript and approved the final version.

